# Correlative microscopy reveals the nanoscale morphology of *E. coli*-derived supported lipid bilayers

**DOI:** 10.1101/2021.11.04.467316

**Authors:** Karan Bali, Zeinab Mohamed, Anna-Maria Pappa, Susan Daniel, Clemens F. Kaminski, Róisín M. Owens, Ioanna Mela

## Abstract

Supported lipid bilayers (SLBs) made from reconstituted lipid vesicles are an important tool in molecular biology. A breakthrough in the field has come with the use of vesicles derived from cell membranes to form SLBs. These new supported bilayers, consisting both of natural and synthetic components, provide a physiologically relevant system on which to study protein-protein interactions as well as protein-ligand interactions and other lipid membrane properties. These complex bilayer systems hold promise but have not yet been fully characterised in terms of their composition, ratio of natural to synthetic component and membrane protein content. Here, we describe a method of correlative atomic force (AFM) with structured illumination microscopy (SIM) for the accurate mapping of complex lipid bilayers that consist of a synthetic fraction and a fraction of lipids derived from *Escherichia coli* outer membrane vesicles (OMVs). We exploit the enhanced resolution and molecular specificity that SIM can offer to identify areas of interest in these bilayers and the atomic scale resolution that the AFM provides to create detailed topography maps of the bilayers. We are thus able to understand the way in which the two different lipid fractions (natural and synthetic) mix within the bilayers, quantify the amount of bacterial membrane incorporated in the bilayer and directly visualise the interaction of these bilayers with bacteria-specific, membrane-binding proteins. Our work sets the foundation for accurately understanding the composition and properties of OMV-derived SLBs and establishes correlative AFM/ SIM as a method for characterising complex systems at the nanoscale.

## Main

Supported lipid bilayers (SLBs) made from reconstituted lipid vesicles are an important tool in molecular biology, especially in the study of biological processes at the cellular or sub-cellular level. However, efforts to deepen our understanding of these processes in physiologically relevant environments are hampered by the simple nature of these bilayers. One approach to introducing physiologically relevant features into SLBs is the reconstitution of purified membrane proteins into proteoliposomes and the subsequent formation of SLBs from these proteoliposomes. This method has been used in several studies including the study of protein-protein interactions^1^, membrane-protein interactions and membrane remodelling^2–4^, membrane poration^5^, host-pathogen interactions^6^ and single receptor activation^7^ among others. Nevertheless, the method of purifying and reconstituting transmembrane proteins requires protein denaturing and refolding in the presence of detergents which leads to low throughput and reproducibility issues. There is also a lack of control over protein orientation in the bilayers.

A breakthrough in the field has come through the use of vesicles derived from cell membranes to form SLBs^8^. For instance, Gram-negative bacteria naturally produce vesicles that contain components of the outer membrane, known as outer membrane vesicles (OMVs). OMVs are 20-250 nm in diameter and are known to function in roles ranging from quorum sensing and signalling to horizontal gene transfer and can be easily isolated and harvested from bacteria through a series of centrifugation steps^9^. By inducing OMVs to rupture and fuse with the addition of synthetic liposomes, it is possible to generate outer membrane SLBs (OM-SLBs) that faithfully represent a naturally occurring outer membrane^10^. These bilayers, which contain both bacterial and synthetic lipid fractions, have been shown to retain components such as outer membrane proteins and lipopolysaccharides in the correct orientation^11^. Therefore, the system retains physiological properties that are beneficial to the investigation of protein-protein interactions, protein-ligand interactions and other lipid membrane properties *in vitro*. These complex bilayer systems hold promise but have not yet been fully characterized in terms of their composition and membrane protein content. Here, we use correlative microscopy to characterise the structural and functional properties of OM-SLBs in previously unseen detail.

To visualise OM-SLBs, we combined atomic force microscopy (AFM) with structured illumination microscopy (SIM). AFM permits the direct observation of SLBs and proteins bound to them at high resolution (∼10 nm)^12^. However, AFM is a label-free technique and is thus incapable of identifying specific proteins of interest on lipid bilayers. Super-resolution microscopy techniques, such as SIM, enable visualisation of specific molecules of interest through staining but at lower resolution than AFM^13^. Here, we combine AFM with SIM to overcome the limitations of the two techniques and visualise OM-SLBs with high resolution and specificity.

To highlight the potential of this approach, we used SLBs that contain OMVs extracted from *E. coli*. We grew *E. coli* BL21(DE3) from an overnight culture and isolated OMVs as described in the Methods section. The dynamic light scattering (DLS) measurements showed an average hydrodynamic size of 101±3 nm, which lies within the size range of OMVs. This was further confirmed by nanoparticle tracking analysis (NTA), a more precise method of size determination compared to DLS since it is able to infer the hydrodynamic radius of particles by resolving their Brownian motion patterns directly^14^. This method identified the main sub peaks at 88 and 152 nm (**Figure S1**). The concentration of vesicles measured with NTA was ∼10^11^ particles/ml.

Transmission electron microscopy was used to directly visualise the OMVs and showed that the vesicles were spherical with diameters in the size range determined by DLS and NTA (Figure S1). Moreover, the vesicles retain the natural composition of the outer membrane, including outer membrane proteins which are involved in a variety of important processes. For instance, we confirm the presence of the membrane protein OmpC, which is involved in the transport of antibiotics and small molecules across as the membrane as well as acting as a binding site for the T4 bacteriophage^15,16^, by dot blot assay (**Figure S2**).

We used these OMVs to form SLBs on coverslips via vesicle fusion, as depicted in **Figure 1.a**. Briefly, in this process the negatively charged OMVs adhere to glass coated with positively charged poly-L-lysine (PLL). The OMVs are induced to rupture and fuse by the addition of palmitoyloleoylphosphatidylglycerol (POPG) liposomes followed by incubation at room temperature for 1 hour. The final stage is the addition of PEG solution, which aids in the bilayer formation process^17^. AFM imaging (**Figure 1.b)** shows the vesicle fusion process taking place, where the approaching edge of the synthetic bilayer induces the rupture of the OMVs. OMVs were stained with the fluorescent lipid-intercalating dye octadecyl rhodamine-18 chloride (R18), and the presence of a complete and mobile bilayer was confirmed by fluorescence recovery after photobleaching (FRAP). The two parameters used to quantify FRAP measurements are the diffusion coefficient (D), which is the mean squared displacement time of the diffusing lipids, and the mobile fraction (MF), which is the proportion of mobile lipids in the bilayer. For the OM-SLB, D and MF values were 0.74±0.14 μm^2^/s and 0.83±0.06, respectively. These values are comparable to those measured in previous studies of SLBs on glass^10^.

**Figure 1:**
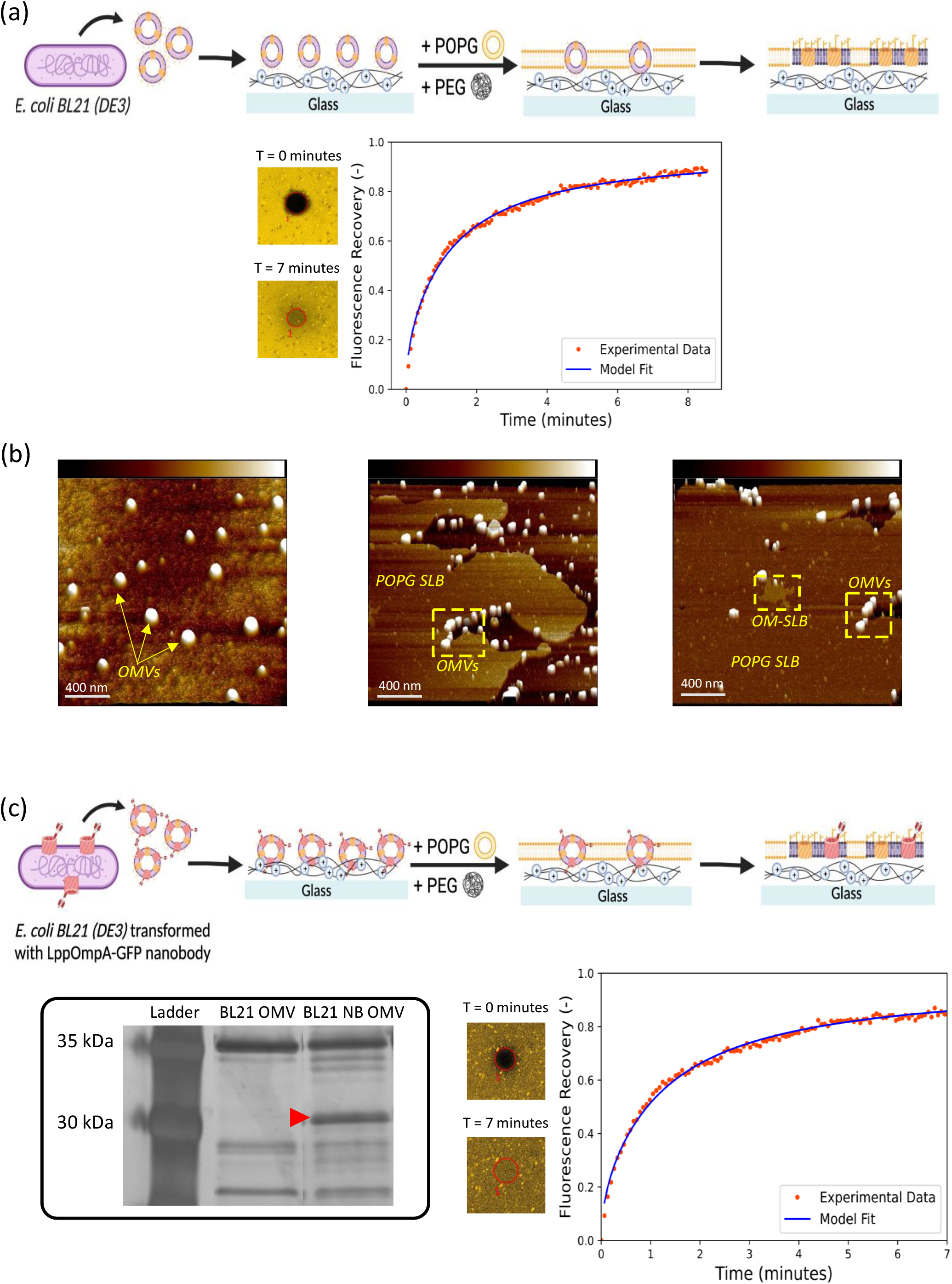
OM-SLB formation and fluorescence microscopy analysis. (a) Schematic showing the process of forming OM-SLBs on PLL-coated cover slips. OMVs are produced naturally by *E. coli* and harvested by ultracentrifugation. The OMVs (∼10^10^ particles/ml) are incubated on the coverslip surface before POPG and PEG are added sequentially to induce rupturing and fusing of the vesicles and the formation of a complete SLB. Naturally occurring membrane proteins are depicted in yellow. Bottom: FRAP data for the OM-SLB, showing that the bleached circle (diameter 30 μm) recovers fluorescence over time; diffusion coefficient (D) and mobile fraction (MF) values are 0.74±0.14 μm^2^/s and 0.83±0.06 respectively. (b) 3D AFM images depicting different stages in the bilayer formation process (note that individual images depict different samples) Left: Intact OMVs on the PLL coated glass surface before rupture. Middle: An area of partially formed bilayer. The synthetic POPG bilayer can be seen on the left side of the image, engulfing the OMVs and causing them to rupture. Right: An area of complete OM-SLB. Height bars are 0-10 nm. (c) Schematic showing the process of forming OM-SLBs, which is the same as (a) except now the OMVs have expressed the nanobody complex (depicted in the schematic in red), via rhamnose induction. Bottom left: SDS-PAGE gel of untransformed and transformed OMVs. The left lane shows the protein content of BL21(DE3) OMVs, whilst the right lane shows the protein content of the LppOmpA-GFP nanobody complex transformed OMVs. A red arrow indicates the nanobody band, which has a molecular weight of ∼28 kDa. Bottom right: FRAP data for the OM-nanobody SLB (diameter of bleached circle 30 μm). The corresponding FRAP parameters are D = 0.94±0.08 μm^2^/s and MF = 0.86±0.10 for this bilayer.

One of the main strengths of the OM-SLB platform is the potential to easily express proteins of interest in the bacteria from which the OMVs are derived, and therefore have these proteins present in the OMVs. We demonstrated this ability by expressing a lipoprotein-outer membrane protein A-GFP binding nanobody (LppOmpA-GFP, hereafter referred to as the nanobody complex) that specifically binds GFP^18^. Using a GFP binding assay, we showed that the nanobody complex was expressed in BL21(DE3) cells (**Figure S3**). We isolated OMVs from the engineered *E. coli* cells as described above, with an extra rhamnose addition step to induce the expression of the nanobody complex, and ran both OMV types on an SDS-PAGE gel to compare the protein content (**Figure 1.c**). An extra protein band was visible in the nanobody OMVs compared with the untransformed OMVs and corresponded to the expected molecular weight of the nanobody (∼28 kDa), indicating successful expression of the protein of interest in the OMVs. By following the same SLB formation sequence as in Figure 1.a, we used the nanobody OMVs to make OM-SLBs containing the nanobody complex. FRAP confirmed the presence of a complete and mobile bilayer with the SLB showing a diffusion coefficient of 0.94±0.08 μm^2^/s and a mobile fraction of 0.86±0.10.

While confocal microscopy and FRAP characterisation of the bilayers indicate the quality of the bilayers and enable quantification of the lipid mobility, the limited resolution of the technique does not allow for precise “mapping” of the synthetic and bacterial components of the OM-SLBs. Such mapping would enable, for example, quantification and optimisation of the ratio of natural to synthetic fraction in the bilayer, quantification of membrane protein content, assessment of the distribution of the bacterial component within the full bilayer, and direct visualisation of membrane-protein interactions. To enable such mapping, we used correlative AFM/SIM (**Figure 2.a**) to sequentially visualize the same area with both techniques, as described in the Methods section.

**Figure 2:**
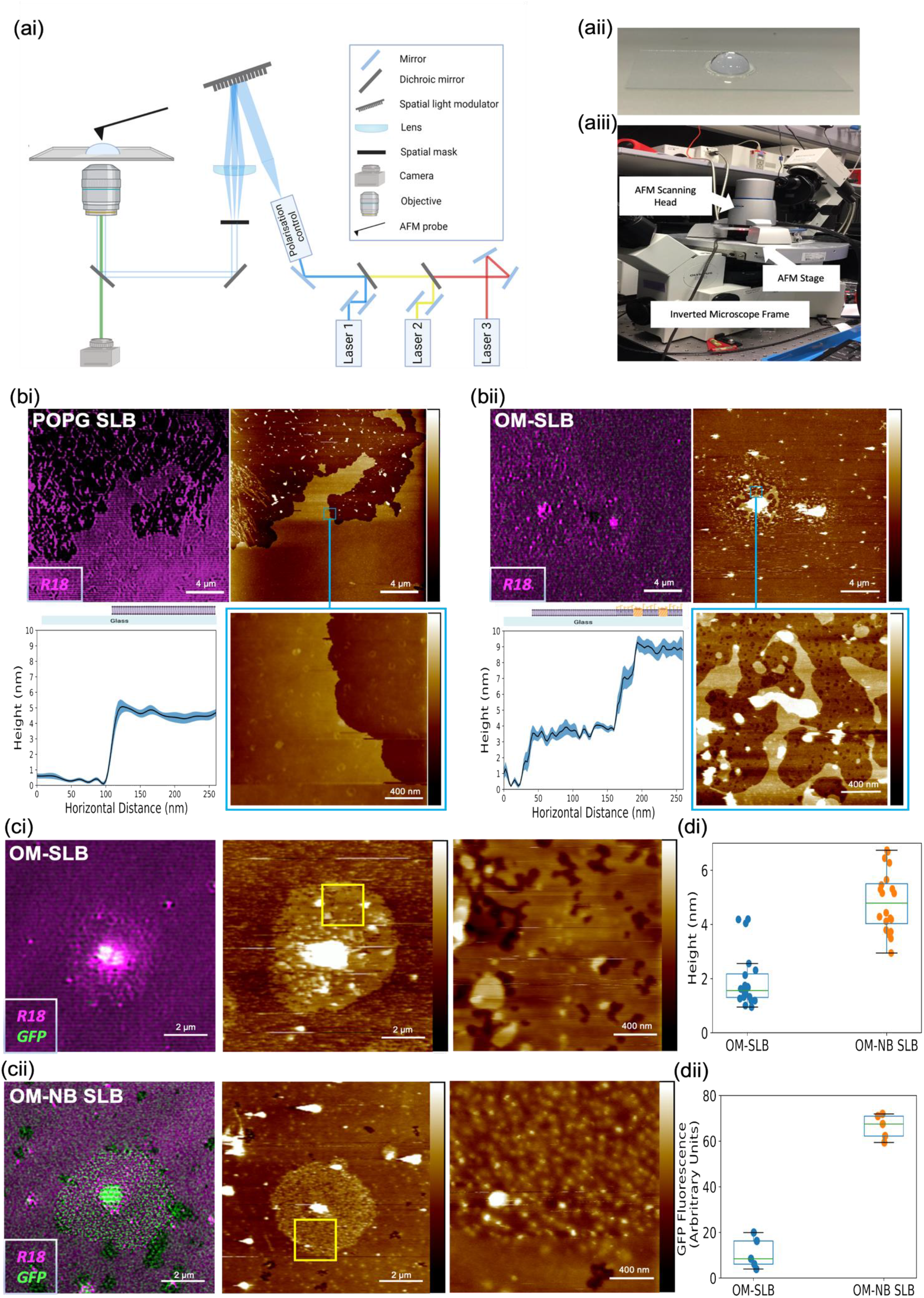
Correlative AFM/SIM analysis of SLBs. (ai) Simplified schematic of the correlative AFM/SIM microscope setup. (aii) Image of a representative sample of an SLB in buffer on a cover slip. (aiii) Annotated image of the correlative microscope setup. (bi) Correlative imaging of a purely synthetic bilayer (POPG-SLB), formed by incubating 4 mg/ml POPG on a PLL-coated cover slip (left). The lipids are stained with R18 and visualised using SIM. (right) The same area is imaged with AFM highlighting the ability of correlative microscopy to access precisely the same area of the bilayer. The inset image (blue box) shows a high magnification area of the bilayer (bottom right). Cross sectional height analysis on this region shows the synthetic SLB to have a height of ∼4 nm. (bii) Correlative imaging of an OM-SLB, formed in the same manner as depicted in Figure 1. The OMVs are stained with R18 and the bilayer is imaged using SIM (left hand image). The same area is imaged with AFM (right hand image). Two distinct regions can be seen in the bilayer: a highly fluorescent region that corresponds to taller features in the AFM image, and one exhibiting lower fluorescence levels where corresponding heights measured by AFM are around 4 nm. High magnification imaging of these areas (iii) shows distinct height regions of bilayer and a cross sectional height analysis shows the two regions of bilayer correspond to heights of ∼ 4 nm and ∼10 nm. These represent the synthetic and bacterial component regions of the bilayer respectively, in line with reported bilayer heights in literature. (c) Comparison of (i) OM-SLB and (ii) OM-SLB expressing the GFP-binding nanobody complex, in both cases after incubation with GFP. (Left to right) Reconstructed SIM image, corresponding AFM image and high magnification AFM image of the yellow box region. (di) Surface roughness analysis for the OM and OM-NB SLBs after GFP incubation. The average height of the surface features in the OM NB SLB case is 4.85 nm, compared to just 1.94 nm for OM SLB case. The height bar for each AFM image is 0-20 nm. (dii) GFP fluorescence intensity signal for the OM-SLB versus the OM-NB SLB, showing the approximately six times increase in fluorescence signal for the nanobody SLB.

We demonstrate the ability of the correlative system to image the same field of view using a POPG-only SLB sample, stained with R18 (**Figure 2.b**). The edge of the bilayer is imaged since the distinctive shape helps to register the SIM and AFM images. Figure 2.b.i shows the same SLB region imaged using SIM and AFM, and by zooming in on an area of the SLB with AFM we obtain a highly precise level of topographical information about the bilayer, showing the bilayer to be smooth with little defects as would be expected from a synthetic bilayer. Furthermore, by taking 5 cross sections of ∼ 250 nm each of the SLB, we show that the average height of the bilayer is 4.4±0.3 nm which corresponds to the height of a synthetic lipid bilayer^19^.

Having established the ability of the correlative microscopy method to track precisely the same area of the bilayer, we move on to imaging SLBs that contain OMVs as depicted in Figure 1. In this case, only the OMVs were stained with R18, while the synthetic POPG lipids were unstained to enable distinction of the two fractions. Upon imaging of the resulting OM-SLB with SIM (**Figure 2.b.ii)**, areas of high and low fluorescence were seen throughout the bilayer, indicating diffusion of phospholipids between the two fractions and demonstrating that the bilayer is complete and continuous. Since the fluorescence from the R18 dye is present in both types of bilayer even though only the OMVs were initially stained, we speculate that this could be attributed to the ability of phospholipids to diffuse between the two bilayers whilst the LPS in the outer leaflet of the OM-SLB diffuses much more slowly, as has been suggested by molecular dynamics simulations^21^. Therefore, areas of high fluorescence were interpreted as originating from OMVs and used as a beacon for the bacterial fraction within the bilayer.

When the same area was imaged using AFM, we saw that the bilayer was not smooth, as was the case for the POPG bilayer, but there were distinct bilayer regions with different heights, which corresponded to areas of low and high fluorescence. Cross sectional height analysis revealed the lower height bilayer region, which corresponded to areas of low fluorescence, to be 3.7±0.5 nm, suggesting this is POPG-SLB. The higher SLB patches, which corresponded to areas of high fluorescence, were 9.1±0.5 nm in height. This height corresponds to the reported height of the OM in *E. coli* cells^22^ which is greater than the height of a synthetic lipid bilayer, owing to the presence of lipopolysaccharides. The combination of SIM and AFM, therefore, reveals that the hybrid SLB contains discrete patches of OM-SLB and POPG-SLB. We quantified the area of the OM-SLB and POPG-SLB patches within the hybrid SLBs in 10 different regions and showed that the bacterial component covers 43 ± 14% of the total bilayer area (**Figure S4**).

Having established that the bacterial membrane forms discrete patches within the artificial SLB, we used correlative AFM/SIM to visualise the localisation of a specific protein within the bacterial component of the hybrid SLBs. We generated standard OM-SLBs and OM-SLBs from bacteria that expressed the nanobody complex and incubated both with GFP. As seen in **Figure 2.b.ii** the OM-SLB shows two SLB regions of distinct height difference. These regions are also evident in the case of the nanobody containing SLB, but crucially here the SIM reconstruction reveals the patches of bacterial SLB fluoresce strongly in the 488 nm wavelength region (**Figure 2.c.ii**) suggesting the presence of bound GFP. This further confirms the existence of bacterial fractions within the bilayer, as the GFP can only bind to the nanobody complex that is present in the OMVs. We quantified the difference in fluorescence between bacterial membrane patches that contain the nanobody complex and those that don’t, by calculating the corrected total green fluorescence (CTF) and showed that the average CTF for the OM-nanobody SLB is approximately six times brighter than that of the OM-SLB (**Figure 2.d.ii**) and that of a POPG SLB (Figure S5). Quantification of surface roughness at the bacterial fractions of the SLBs showed that the mean height of the surface features in the nanobody-SLB with GFP bound was 4.85 nm (range 2.95–6.74 nm). By contrast, surface features in the control OM-SLB had a mean height of 1.94 nm (range 0.94–4.20 nm). This shows that the presence of the bound GFP increases the roughness of the surface. Although there are outlier points in the OM-SLB roughness analysis, these likely reflect the presence of naturally occurring outer membrane proteins. The ability to map the bacterial component of the SLB using GFP binding is a key finding, as it shows that not only can we identify areas of interest in these complex systems using correlative AFM-SIM, but we can also quantify binding events occurring on these membranes. Furthermore, we can manipulate these systems precisely by altering the expression profile of the bacterial component. This is particularly exciting when we consider the future applications of this method, particularly with respect to antimicrobial screening studies^11^.

In conclusion, we present here correlative AFM/SIM as a method for accurate characterisation of bacterial - derived supported lipid bilayers at the nanoscale. This approach enables not only the mapping and quantification of the bacterial and synthetic components within those bilayers, but also the visualisation of single proteins bound to those components. Having access to detailed maps of the bacterial and synthetic components of these bilayers enables optimisation of these systems with respect to the quality of the bilayers and the quantity of the bacterial fraction. Our method has the potential to facilitate antimicrobial discovery by enabling investigation of how antimicrobial drugs and drug delivery vehicles, such as nanomaterials and bacteriophages, interact with the bacterial membrane. In the present study, we have optimised our method for visualization of SLBs that contain bacterial membrane fractions, but correlative AFM-SIM can be employed to characterise SLBs that contain natural components derived from mammalian cells, plants and other organisms. Furthermore, the method can be adapted for a variety of other applications, ranging from investigation of single-molecule interactions to characterization of inorganic 2D materials.

## Methods

### Growth of bacterial cultures and isolation of OMVs

5 ml of liquid Lura-Bertani (LB) broth was inoculated with *E. coli BL21 (DE3)* (Invitrogen) cells for 16-20 hours to make an overnight culture. 2 ml of the overnight culture was added to 200 ml of LB broth and allowed to incubate at 37°C for ∼3 hours until the OD600 of the culture was ∼1.5. The culture was then centrifuged (4000xg, 4C) for 15 minutes in order to remove cell debris, and the supernatant was collected. The supernatant was further passed through a 0.22 μm filter. The outer membrane vesicles (OMVs) were then isolated by ultracentrifugation (140 000xg, 4°C) for 3 hours (Beckman Coulter, Type 50.2 Ti Fixed-Angle Rotor) and the pellets were resuspended in 250 μl of Phosphate Buffered Saline (PBS) supplemented with 2 mM MgCl_2_ solution. Finally, the OMV solution was centrifuged (16 000xg, 4°C) for 30 minutes to remove any final contaminants such as flagella. The supernatant was collected and re-suspended in 500 μl of PBS supplemented with 2 mM MgCl_2_ solution. The final OMV solutions were then stored at -80°C for further experiments.

### Preparation of synthetic liposomes

1-palmitoyl-2-oleoyl-sn-glycero-3-phospho-(1’-rac-glycerol) (POPG), purchased from Avanti Polar Lipids and stored in chloroform solution at -20°C, was used to prepare synthetic lipid liposomes. A N_2_ stream was used to evaporate the chloroform, and the sample was further desiccated for 1 hour in a vacuum. The lipids were then hydrated in PBS supplemented with 2 mM MgCl2 to give a final lipid concentration of 4 mg/ml. Single unilamellar vesicles were made by lipid extrusion through a 50 nm pore sized polycarbonate membrane, and samples were stored for up to two weeks at 4oC.

### Formation of supported lipid bilayers on glass coverslips

Glass coverslips (Academy, 22×40 mm, 0.16-0.19 mm thick) were first cleaned with Acetone and Isopropanol before being functionalised by incubating with poly-l-lysine solution (0.1% w/v) for 15 minutes. The PLL solution was washed away with DI H_2_O before 100 μl of ∼10^10^ OMVs/ml was added to the glass slide. The OMVs were allowed to incubate for 20 minutes before washing twice with PBS solution in order to remove excess unadhered OMVs. 100 μl of the synthetic lipid vesicles were then added for 1 hour in order to induce rupturing of the OMVs. The well was then washed again twice with PBS and finally 30% PEG was added for 10 minutes as a final SLB formation step. The SLBs were then kept in PBS solution for imaging. For experiments involving only synthetic lipid bilayers, the same protocol was followed without the first OMV incubation step.

### Characterisation of OMVs

#### 1. Dynamic Light Scattering

Dynamic Light Scattering (DLS) measurements were performed using a Zetasizer Nano S90 (Malvern Panalytical) configured with a 633 nm laser and a 90o scattering optic. 1 ml of sample was transferred into a disposable plastic cuvette, and three runs were taken for each measurement. The intensity of the scattered light is used by the Zetasizer software to determine three main parameters: Z average (d.nm) which measures the average size of the particle distribution; count rate (kcps) which counts the number of photons detected per second and is related to the concentration and quality of the sample; polydispersity index (PDI) which provides a measure for the heterogeneity of the particle size distribution.

#### 2. Nanoparticle Tracking Analysis

Nanoparticle Tracking Analysis (NTA) was carried out at the University of Cambridge Veterinary School. Samples were analysed using a Nanosight NS500 (Malvern Panalytical) fitted with an Electron Multiplying Charged Couple Device (EMCCD) camera configured with a 522 nm laser. Prior to analysis, samples were diluted (1:500) in PBS. 5 × 60 second videos were recorded for each sample analysed, with a temperature range of 20.8 – 21.5oC and a camera level of 15. NTA 3.2 software was used to analyse the data with a detection threshold of 5.

#### 3. Transmission Electron Microscopy

Transmission Electron Microscopy (TEM) was carried out at the Cambridge Advanced Imaging Centre. 10 μl of sample was negatively stained with 1% (w/v) uranyl acetate solution for 2 minutes at room temperature before being visualised with a Tecnai G2 80-200 keV transmission electron microscope, operating at 200 keV with images recorded with a bottom-mounted AMT CCD camera.

### Characterisation of SLBs using fluorescence recovery after photobleaching (FRAP)

Prior to analysis by fluorescence recovery after photobleaching (FRAP), OMVs must be fluorescently labelled. This was achieved by adding 1 μl of octadecyl rhodamine chloride 18 dye (Invitrogen) to 200 μl of OMV solution, and sonicating for 15 minutes. A G25 spin column (GE healthcare) was used to remove unbound/excess R18 by centrifugation at 3000 rpm for 3 minutes at room temperature. Lipid bilayer formation was then conducted using the protocol outlined above.

FRAP measurements were conducted using an inverted Zeiss LSM800 confocal microscope with a 10x objective lens. A 30 μm diameter bleaching spot was made, and recovery of the fluorescence intensity of this spot was measured over time relative to a 50 μm diameter reference spot. The data were analysed using MATLAB, and a Soumpasis fit was made in order to extract the diffusion coefficient (D) according to the equation D = r^2^/4 where is the radius of the photobleached spot and τ is the characteristic diffusion time.

### Production of OMVs expressing Lpp-OmpA-GFP binding nanobody

The plasmid pK:LppOmpA-NB was kindly provided by Dr Morten Norholm and transformed into *E. coli* BL21 (DE3) cells as described previously^18^.In order to express the protein in OMVs, OMVs were grown in the same manner as described above except that 5 mM rhamnose was added to the bacteria culture at OD ∼0.3 to induce nanobody expression. The culture was then grown for a further 5 hours after induction to ensure nanobody OMV production.

### GFP binding assays for E. coli BL21 cells

Cells were pelleted from an overnight culture via centrifugation for 4 minutes at 2272xg, and resuspended in 50 μl 50 mM Tris buffer, before being mixed with 50 μl 0.12 mg/ml GFP solution (Sino Biological) and incubated for 30 minutes at 30°C at 250-300 rpm. The incubation was stopped by centrifugating the cells for 4 minutes at 2272xg, and then the cells were washed twice with 300 μl 50 mM Tris buffer and resuspended in 200 μl 50 mM Tris buffer for imaging.

### GFP binding assays for OM SLBs

Bilayers were formed by the process described previously and incubated with 0.06 mg/ml GFP solution (Sino Biological) for 30 minutes at 30°C. The incubation was stopped by removing the GFP solution and washing with PBS buffer three times before imaging.

### SDS-PAGE analysis of OMVs

OMVs were boiled at 95°C for 5 minutes with lithium dodecyl sulfate (LDS) sample buffer (4X Bolt; Invitrogen) and run on a NuPAGE 12% Bis-Tris gel (Invitrogen) in MOPS buffer at 200 V for 30 minutes. The protein bands were visualised using a ProteoSilver Silver Stain kit (Sigma-Aldrich).

### Dot Blot against OmpC

OMVs and POPG (used as a negative control) were targeted with OmpC monoclonal antibody (Bioorbyt). Goat Anti-Mouse IgG/ Horseradish Peroxidase Conjugate (HRP) (Novex, Life Technologies) was used as the secondary antibody. A PVDF membrane (ThermoFisher Scientific) with 0.45 µm pore size was activated by shortly immersing it in methanol, before being incubated in transfer buffer (Novex, Life Technologies) at room temperature for 3 minutes. The membrane was transferred onto a filter paper (ThermoFisher Scientific), previously soaked in transfer buffer, and 10 µL of each sample were added onto the membrane. The membrane was incubated in blocking buffer (0.1 % Tween 20 (ThermoFisher Scientific) and 5 % BSA (ThermoFisher Scientific) in PBS) at room temperature for 1hour to block unspecific sites, then it was incubated in primary antibody solution (1:1000 dilution in blocking buffer) at room temperature for 1 hour. The membrane was washed three times in washing buffer (0.1 % Tween 20 in PBS) at room temperature for 2 minutes, then incubated with secondary antibody solution (1:1000 dilution in washing buffer) at room temperature for 1 hour. The membrane was washed three times in washing buffer at room temperature for 2 minutes, before being transferred to a transparent film. After addition of the chemiluminescent solution (SuperSignal West Pico PLUS, ThermoFisher), the membrane was transferred to a black background for imaging (G:BOX mini 6/9, Syngene).

### Correlative AFM/SIM and data analysis

Correlative AFM–SIM imaging was performed by combining a Bioscope Resolve system (Bruker) with a custom-built SIM system^23^. The piezo stage of the SIM microscope was removed from the inverted microscope frame and the stage of the AFM system was used to drive both microscopes at the same time. The stage of the specific AFM system is designed so that the sample holder allows for optical detection of specimens from below, while the AFM scanning head can access the sample from above. The fields of view (FOVs) of the two microscopes were aligned so that the AFM probe was positioned in the middle of the FOV of the SIM microscope, by carefully moving the AFM stage using the alignment knobs. The final, fine alignment was achieved by using a bright-field image of the ‘shadow’ of the AFM cantilever, taken with the SIM, to precisely align the AFM probe with the SIM lens (Supplementary Figure 6).

To acquire structured illumination microscopy images, a ×60/1.2 NA water immersion lens (UPLSAPO 60XW, Olympus) focused the structured illumination pattern onto the sample, and the same lens was also used to capture the fluorescence emission light before imaging onto an sCMOS camera (C11440, Hamamatsu). The wavelengths used for excitation were 561 nm (OBIS 561, Coherent) for the lipid bilayers and 488 nm (iBEAM-SMART-488, Toptica) for the GFP. Images were acquired using customized SIM software described previously^23^.

AFM images were acquired in Scanasyst mode using ScanasystFluid+ probes (Bruker), with a nominal spring constant of 0.7 N m-1 and a resonant frequency of 150 kHz. Images were recorded at scan speeds of 1.5 Hz and tip–sample interaction forces between 200 and 300 pN. Large–scale images (20 × 20 μm) were used to register the AFM with the SIM FOVs and small (2 × 2 μm) scans were performed to better resolve the morphology of the bilayers. Raw AFM images were first order fitted with reference to the lipid bilayer. Height measurements on the bilayers were performed by taking cross-sections across different areas of interest, using the Nanoscope analysis software (Bruker).

For quantification of the bacterial component in the bilayers, AFM micrographs were converted to 8-bit images using Fiji (ImageJ) and thresholded to the height of the synthetic bilayer component. The area covered by bacterial component (above the threshold) was calculated using the inbuilt area measurement tool in Fiji.

## Acknowledgements

The authors thank Dr Morten Nørholm (Novo Nordisk Foundation Center for Biosustainability, Technical University of Denmark) for kindly providing the LppOmpA-GFP binding nanobody plasmid and Dr Emmanuel Derivery (MRC-LMB, Cambridge) for useful discussions. KB acknowledges funding from Engineering and Physical Sciences Research Council Doctoral Training Program (EPSRC DTP). C.F.K. acknowledges funding from the Engineering and Physical Sciences Research Council, EPSRC (EP/ H018301/1, EP/L015889/1); Wellcome Trust (089703/Z/ 09/Z, 3-3249/Z/16/Z); Medical Research Council MRC (MR/K015850/1, MR/K02292X/1); MedImmune; and Infinitus (China), Ltd. R.M.O. acknowledges funding for this project, sponsored by the Defense Advanced Research Projects Agency (DARPA) Army Research Office and accomplished under Cooperative Agreement Number W911NF-18-2-0152. The views and conclusions contained in this document are those of the authors and should not be interpreted as representing the official policies, either expressed or implied, of DARPA or the Army Research Office or the U.S. Government. The U.S. Government is authorized to reproduce and distribute reprints for Government purposes notwithstanding any copyright notation herein. I.M. acknowledges funding from the National Biofilms Innovation Centre (BB/R012415/1 03PoC20-105).

