## Supplementary Data for "Correlative microscopy reveals the nanoscale morphology of *E. coli*-derived supported lipid bilayers"

### Supplementary Material

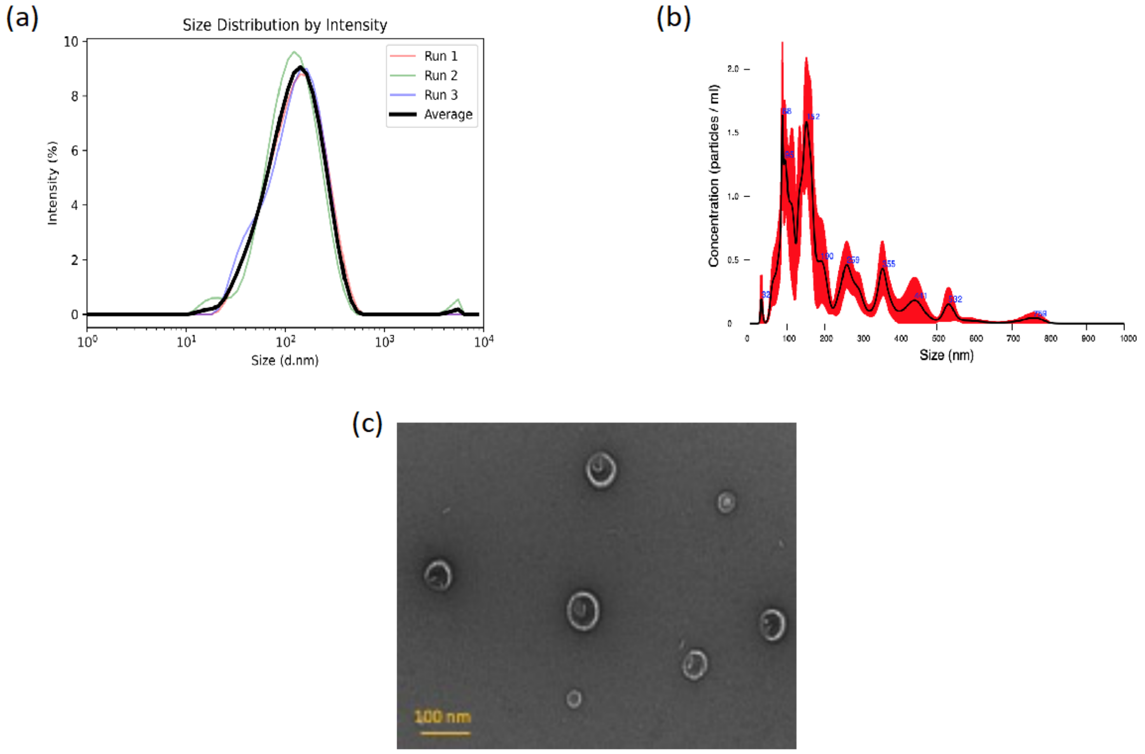


Figure S1: OMV characterisation. (a) DLS shows that the size of OMVs is in the expected 20-200 nm range (b) NTA shows more precise size mapping, with the major OMV size peaks being at 88,95 and 152 nm (c) TEM image showing the OMVs.

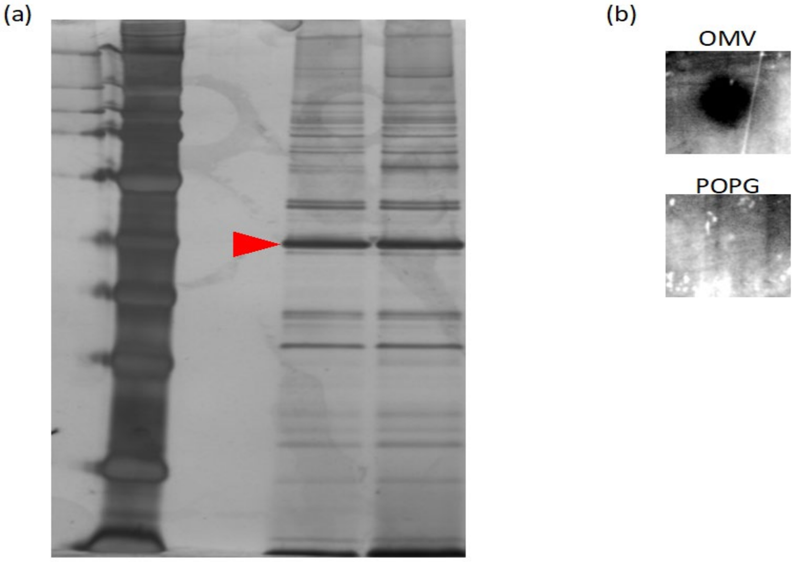


Figure S2: Characterisation of protein content of BL21 OMVs. (a) SDS-PAGE gel for two independently isolated BL21 OMV samples. A prominent band can be seen at ~40

kDa, which corresponds to the molecular weight of OmpC. (b) Dot blots of OMV and POPG samples against an OmpC marker. In the OMV sample, a dark spot can be clearly seen due to the secondary HRP antibody attaching to the OmpC primary antibody that has bound to OmpC proteins in the OMVs. The POPG sample shows no signal since the OmpC primary antibody is unable to bind.

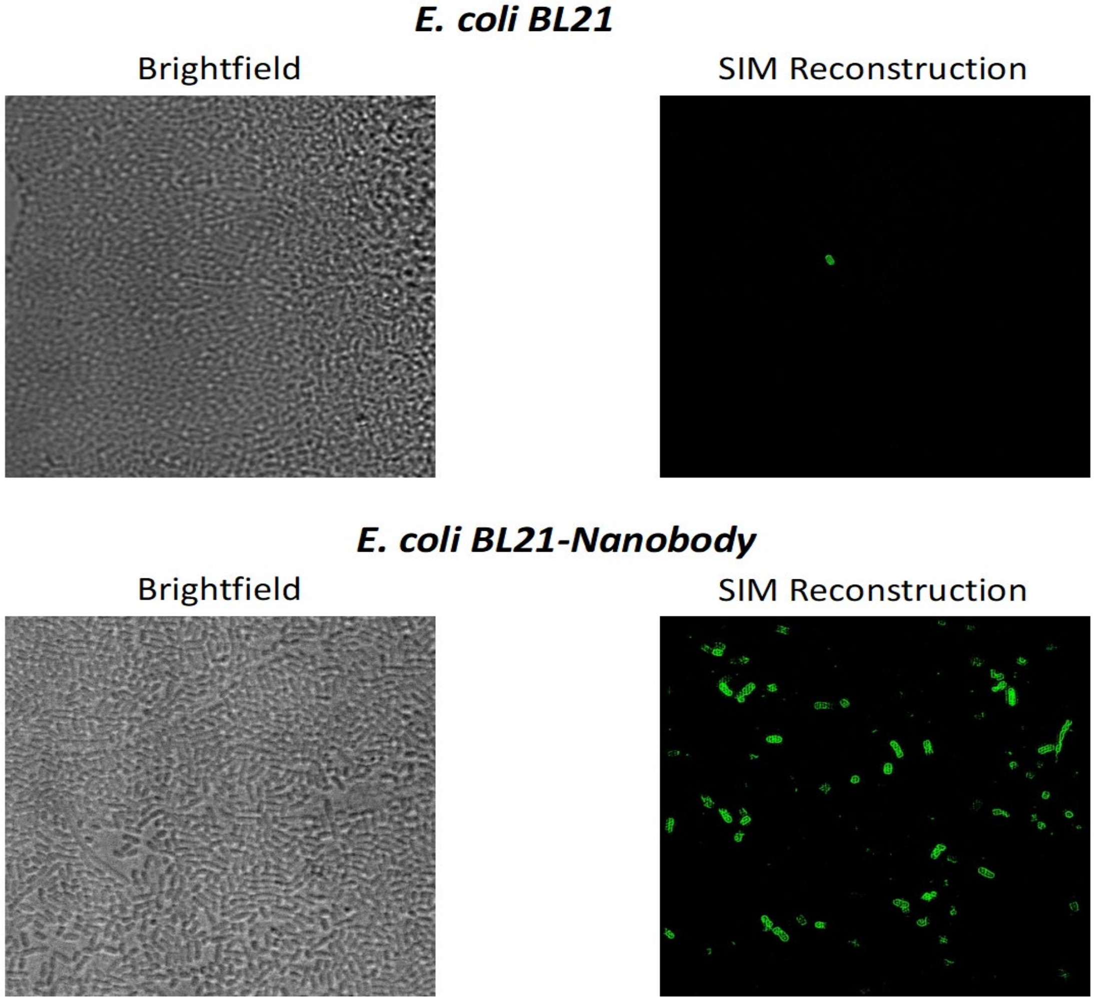


Figure S3: Whole cell *E. coli* transformed with LppOmpA-nanobody binding GFP assay. 0.01 mg/ml GFP solution is incubated with cells for 20 minutes at 30ºC, after which the cells are centrifuged, resuspended and washed in Tris buffer before imaging. In the BL21 sample, minimal interaction between the bacteria and GFP can be observed, while the BL21-nanobody sample shows a large number of bacteria decorated with GFP. Crucially, the signal arises from the surface of the cells, suggesting the GFP is binding to a surface protein as opposed to being internalised.

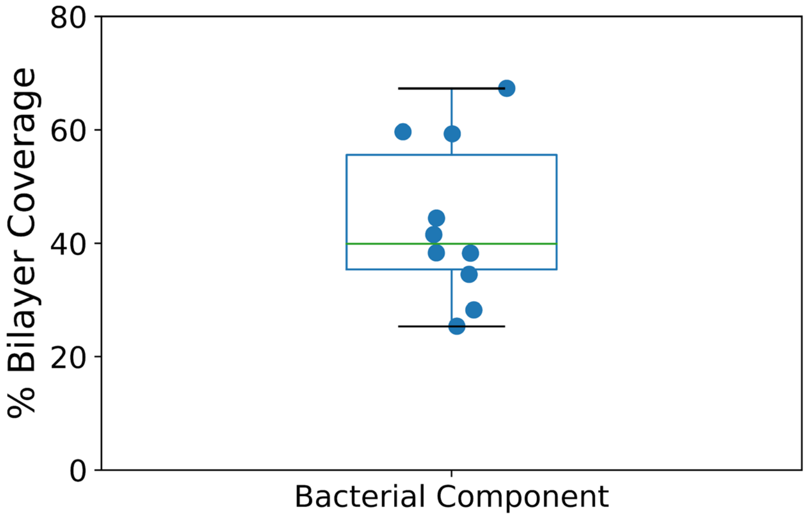


Figure S4: Quantification of the bacterial fraction within the OM-SLBs.

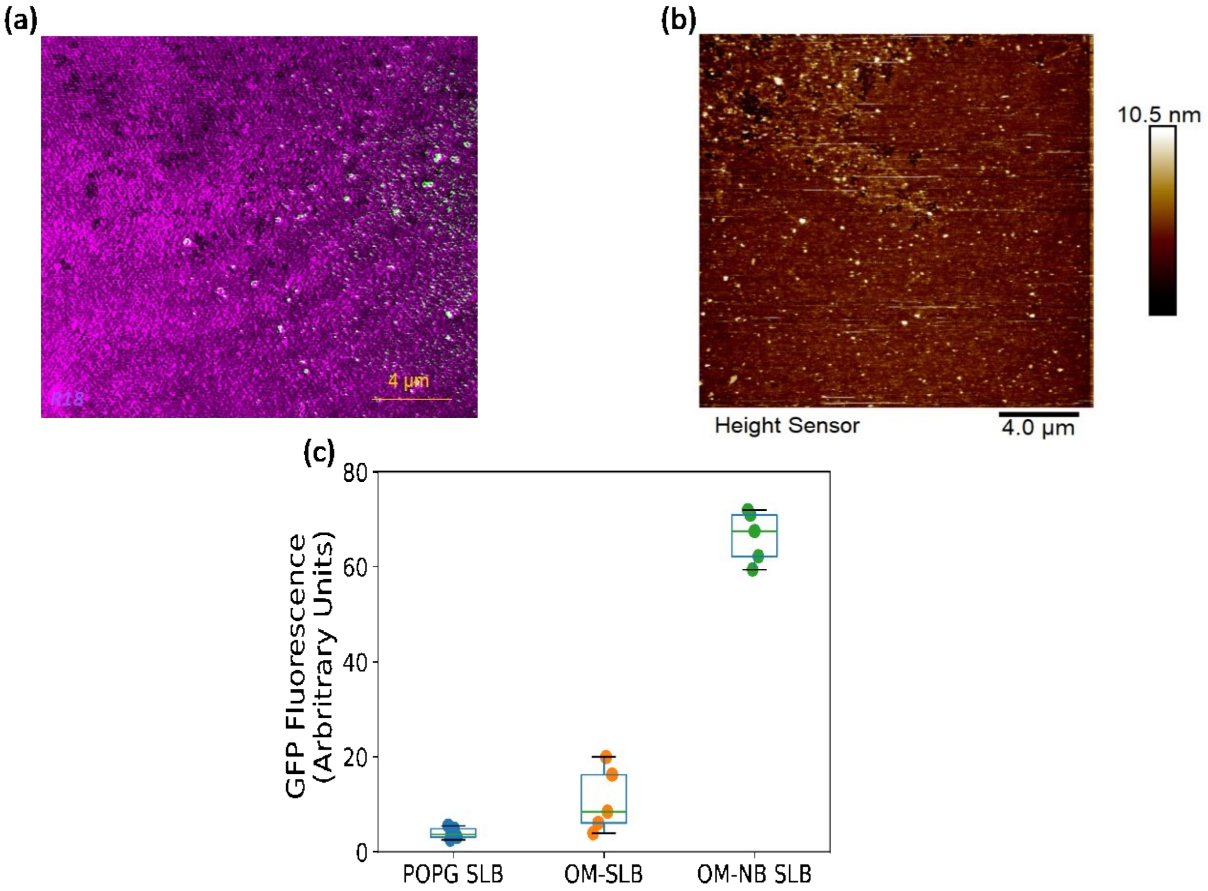


Figure S5: Correlative AFM/SIM images for POPG SLB incubated with GFP. (a) Reconstructed SIM image (b) AFM image. In both cases there is negligible GFP signal or binding. (c) Corrected total fluorescence (CTF) in the 488 nm range for the bacterial component region of OM-SLB vs OM-NB SLB vs representative region of POPG SLB. The bar chart clearly shows the increase fluorescence in the nanobody case compared to the two controls.

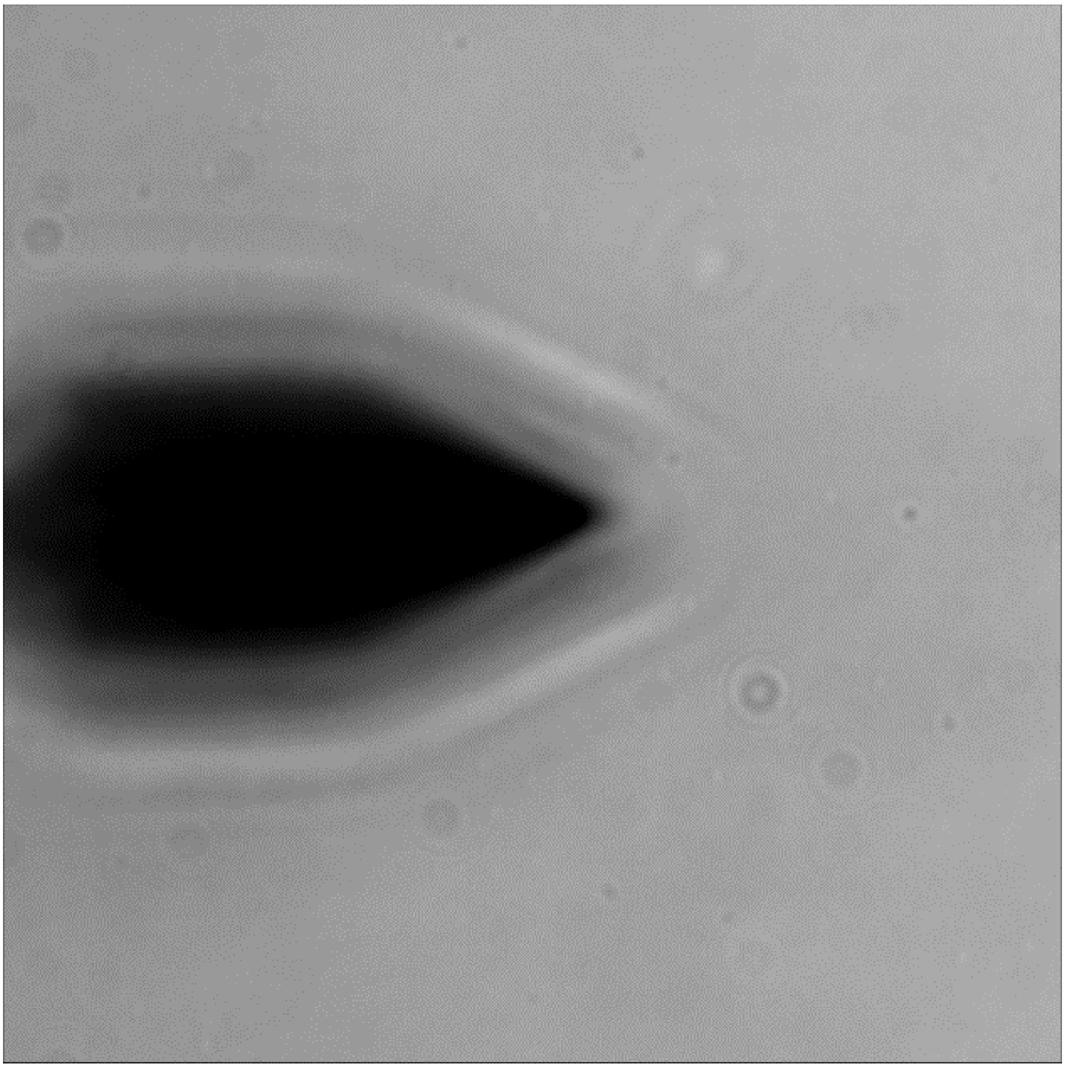

Figure S6: The AFM cantilever as imaged through the SIM microscope. The tip of the cantilever is positioned in the middle of the field of view of the SIM, ensuring the accuracy of the alignment of the fields of view of the two microscopes.
